# Prenatal SARS-CoV-2 infection alters postpartum human milk-derived extracellular vesicles

**DOI:** 10.1101/2023.06.01.543234

**Authors:** Somchai Chutipongtanate, Hatice Cetinkaya, Xiang Zhang, Damaris Kuhnell, Desirée Benefield, Wendy Haffey, Michael Wyder, Richa Patel, Shannon C. Conrey, Allison R. Burrell, Scott Langevin, Laurie Nommsen-Rivers, David S. Newburg, Kenneth D. Greis, Mary A. Staat, Ardythe L. Morrow

**Affiliations:** Department of Environmental and Public Health Sciences, University of Cincinnati College of Medicine, Cincinnati, OH 45267 USA; Department of Molecular Cellular Biosciences, Center for Advanced Structural Biology, University of Cincinnati College of Medicine, Cincinnati, OH 45267 USA; Department of Cancer Biology, University of Cincinnati College of Medicine, Cincinnati, OH 45267 USA; Department of Infectious Disease, Cincinnati Children’s Hospital Medical Center, Cincinnati, OH 45267 USA; Department of Rehabilitation, Exercise, and Nutrition, University of Cincinnati College of Allied Health Sciences, Cincinnati, OH 45267 USA

**Keywords:** Breast milk, COVID-19, Exosomes, MicroRNAs, Proteomics, Surfaceomics

## Abstract

Human milk-derived extracellular vesicles (HMEVs) are crucial functional components in breast milk, contributing to infant health and development. Maternal conditions could affect HMEV cargos; however, the impact of SARS-CoV-2 infection on HMEVs remains unknown. This study evaluated the influence of SARS-CoV-2 infection during pregnancy on postpartum HMEV molecules. Milk samples (9 prenatal SARS-CoV-2 vs. 9 controls) were retrieved from the IMPRINT birth cohort. After defatting and casein micelle disaggregation, 1 mL milk was subjected to a sequential process of centrifugation, ultrafiltration, and qEV-size exclusion chromatography. Particle and protein characterizations were performed following the MISEV2018 guidelines. EV lysates were analyzed through proteomics and miRNA sequencing, while the intact EVs were biotinylated for surfaceomic analysis. Multi-Omics was employed to predict HMEV functions associated with prenatal SARS-CoV-2 infection. Demographic data between the prenatal SARS-CoV-2 and control groups were similar. The median duration from maternal SARS-CoV-2 test positivity to milk collection was 3 months (range: 1-6 months). Transmission electron microscopy showed the cup-shaped nanoparticles. Nanoparticle tracking analysis demonstrated particle diameters of <200 nm and yields of >1e11 particles from 1 mL milk. Western immunoblots detected ALIX, CD9 and HSP70, supporting the presence of HMEVs in the isolates. Thousands of HMEV cargos and hundreds of surface proteins were identified and compared. Multi-Omics predicted that mothers with prenatal SARS-CoV-2 infection produced HMEVs with enhanced functionalities involving metabolic reprogramming and mucosal tissue development, while mitigating inflammation and lower EV transmigration potential. Our findings suggest that SARS-CoV-2 infection during pregnancy boosts mucosal site-specific functions of HMEVs, potentially protecting infants against viral infections. Further prospective studies should be pursued to reevaluate the short- and long-term benefits of breastfeeding in the post-COVID era.

## INTRODUCTION

Human milk, a complex and dynamic biofluid, provides nutrients that support infant growth and bioactive components which protect infants against various diseases,^1,2^ including respiratory infections in early and late childhood.^3^ Clinical trials and epidemiologic studies confirm the beneficial effects of feeding human milk over infant formula in preventing early and long-term diseases.^2-4^ Cumulative evidence of the mechanisms show breastfeeding delivers positive health outcomes to children through human milk bioactive components, including immunoglobulins, growth factors, cytokines, adipokines, hormones, lipids, peptides, non-digestible oligosaccharides, cells and extracellular vesicles (EVs).^5-11^

Human milk-derived EVs (HMEVs) are lipid bilayer-enclosed nanoscale vesicles mainly secreted by mammary epithelial cells and play roles in physiological functions and pathological processes.^11-13^ HMEVs carry selective molecular cargos from mammary glands with the potential to modulate gene expression and cell signaling in infant tissues.^11-13^ Current evidence suggest that HMEVs represent a vital mechanism of maternal-to-child intergenerational communication.^11-13^ The infant intestinal mucosa appears to be a primary target of HMEVs. Appreciable amounts of HMEVs are absorbed from the gastrointestinal tract into blood circulation, and from there can reach various organs to modulate functions in recipient cells in vitro and in vivo.^14-17^ Studies have shown that maternal pathological conditions during pregnancy affect HMEV molecular cargos with potential functional changes in breastfed infants,^11,18-20^ while the exact mechanisms remain to be elucidated.

The coronavirus disease 2019 (COVID-19) pandemic, caused by severe acute respiratory syndrome coronavirus-2 (SARS-CoV-2) infection, presents a significant challenge to public health globally. While COVID-19 has had substantial effects on breastfeeding practices around the world due to concerns about possible mother-to-infant transmission,^21^ there is no evidence of direct viral transmission through breastfeeding.^22,23^ Exposure to SARS-CoV-2 has been shown to influence human milk components, including antibodies, inflammatory mediators, cytokines and immune cells.^24-33^ Nevertheless, our knowledge about the impact of maternal SARS-CoV-2 infection during pregnancy on HMEVs remains limited.

The NIH-funded IMPRINT birth cohort has implemented a standardized protocol for the collection of breast milk at 2 weeks, 6 weeks, and annually during the summer (ranging from 9 to 52 weeks). During the COVID-19 pandemic, a substantial proportion of participating mothers experienced SARS-CoV-2 infection during pregnancy, thereby providing an unparalleled opportunity to examine the potential effects of prenatal SARS-CoV-2 infections on the constituents of postpartum human milk, inclusive of HMEVs.

In close collaboration with the IMPRINT cohort, the aim of this study was to ascertain the influence of prenatal SARS-CoV-2 infection on HMEV molecular cargos. Following the optimization of our previously established method^34-36^ for the successful isolation of EVs from the complex matrix of human milk, we performed a multi-Omic analysis. This incorporated proteomics, surfaceomics (or the proteomic analysis of surface EV proteins), and miRNA sequencing on HMEVs isolated from 9 mothers with prenatal SARS-CoV-2 infection, compared against 9 healthy control subjects. Alterations in the HMEV cargos were detected and functional predictions were carried out based on those changes to elucidate potential impacts on child health outcomes. Our findings suggest that SARS-CoV-2 infection during pregnancy may boost mucosal site-specific functions of HMEVs in breastfed infants.

## MATERIALS AND METHODS

### HMEV isolation

Human milk samples (1 mL/sample) were retrieved from the IMPRINT birth cohort (protocol ID 2019-0629). Milk samples were subjected to stepwise centrifugation (500 x g for 15 min, 3000 x g for 15 min) to remove cells and milk fat, added sodium citrate to the final concentration of 1% (w/v) and incubated at 4°C for 30 min to disrupt casein micelles, and centrifuge at 14,000 x g for 30 min to eliminate microvesicles. The supernatant was then concentrated using a 100-kDa cutoff centrifugal filter (Thermo Scientific) to achieve 0.5 mL. The concentrated supernatant was loaded onto the Izon qEV size-exclusion column (qEVoriginal/35nm) (Izon Science, UK) to isolate the EV particles out of soluble protein contaminates. The particle eluates were pooled and concentrated by 100-kDa cutoff centrifugal filtration (Thermo Scientific), resulting in the final volume of 1 mL. Pierce660 assay (Thermo) was used to estimate EV protein amounts. All isolation procedures were performed at 4°C, or on ice, as appropriate. The isolated EVs were aliquoted (100 μL per aliquot) and kept at -80 °C until used.

### Nanoparticle tracking analysis (NTA)

NanoSight NS300 (Malvern Instruments Ltd., Malvern, Worcestershire, UK) was used for particle size distribution and concentration analysis. The sample was diluted 1:1000 in PBS to a final volume of 1 mL. The diluted sample was injected using a 1-mL syringe. Five 1-min videos were captured for each sample with the following parameters; camera: sCMOS; cell temperature: 25°C; After capture, the videos were analyzed by NanoSight Software NTA 3.4 Build 3.4.003 with this setting; detection threshold: 5; blur size and max jump distance: auto. Ideal concentrations were measured at 20-100 particles/frame.

### Transmission electron microscope

Five microliters of the EV sample was loaded onto a 200 mesh grid (FCF200-CU; Electron Microscopy Sciences) and negatively stained by using 2% uranyl acetate (#22400; Electron Microscopy Sciences) in 50% methanol for 1 min and then dried at room temperature. Imaging was performed using a transmission electron microscope (Thermo Talos L120C Transmission Electron Microscope) with an acceleration voltage of 100 kV.

### Western blot analysis

The EV sample (10 μg protein) was mixed with Leammli buffer, heated at 95°C for 10 min, resolved in a 4-12% Invitrogen B-T gel using MOPS buffer and transferred onto a polyvinylidene difluoride (PVDF) membrane using BioRad transblot (Bio-Rad, Hercules, CA, USA). The membrane was blocked with 5% skim milk, and then probed with primary antibodies as follows; anti-Hsp70 (1:1000) (#sc-373867; Santa Cruz Biotechnology, Dallas, TX), anti-CD9 (1:1000) (#sc-13118; Santa Cruz Biotechnology), anti-Alix (1:1000) (#2171; Cell Signaling Technology, Inc., Danvers, MA) antibodies for at 4°C overnight. After washing, the membranes were incubated with anti-mouse IgG-HRP (1:3000) (#NA931, MilliporeSigma) at room temperature for 1 h. The immunoblot was developed by SuperSignal West Pico PLUS Chemiluminescent (ThermoFisher), and imaged on a ChemiDoc Touch Imaging system (BioRad Laboratories, Inc., Hercules, CA).

### NanoLC-MS/MS

Two micrograms of EV proteins were dried in a speedvac and resuspended in 15 μL of 50 mM ammonium bicarbonate. Samples were then reduced by dithiothreitol (DTT; a final concentration of 10 mM) with heat at 95°C for 5 min, alkalinized by iodoacetamide (IAA; a final concentration of 10 mM) for 20 min at room temperature in dark, and digest by trypsin (1:50 w/w; MS-grade, Thermo) at 37°C overnight. Samples were desalted by C18 solid phase extraction, dried by a speedvac, and resuspended in 0.1% formic acid in H_2_O. Data were collected on an Orbitrap Eclipse mass spectrometer coupled to Dionex Ultimate 3000 RSLCnano (Thermo Fisher Scientific). Peptide was injected onto a 5-mm nanoViper μ-Precolumn (internal diameter [i.d.] 300 mm, C18 PepMap100, 5.0 μm, 100 Å; Thermo Fisher Scientific) at 5 μl/min in 0.1% formic acid in H_2_O for 5 min. For chromatographic separation, the trap column was switched to align with EASY-Spray column PepMap RSLC C18 with a 150-mm column (i.d. 75 μm, C18, 3.0 μm, 100 Å). Peptides were eluted using variable mobile phase gradient from 98% phase A (0.1% formic acid in H_2_O) to 32% phase B (0.1% formic acid in acetonitrile [ACN]) for 60 min at 300 nl/min. MS1 was collected in Orbitrap (120,000 resolution; maximum injection 50 ms; automatic gain control [AGC] 4×10^5^). Charge states between 2 and 6 were required for MS^2^ analysis with a 20-s dynamic exclusion window and cycle time of 2.5 s. MS^2^ scans were performed in an ion trap with higher-energy collisional dissociation (HCD) fragmentation (isolation window 0.8 Da; normalized collision energy (NCE) 30%; maximum injection 40 ms; AGC 5×10^4^) and recorded using Xcalibur 4.3 (Thermo Fisher Scientific). The raw files were searched using Proteome Discoverer v.2.4 (Thermo Fisher Scientific) against human protein database and the Sequest HT search algorithm with LFQ parameter to identified and quantified peptides at FDR<1% before protein inference (reported at >99% confidence). The results were exported as the Excel format for further analysis. Raw expression data was log2 transformed, normalized to reduce technical variations, and imputed for missing values before differential expression analysis and functional prediction.

### Surfaceomic analysis

EV surface protein enrichment was performed by Pierce cell surface protein biotinylation and isolation kit (Thermo Scientific) as previous study^37^ with a modification on washing procedure (replacing ultracentrifugation by ultrafiltration). Nine intact EV samples generated three pooled EVs (equivalent to 200 μg EV proteins/pool). The pooled EV samples were labeled with sulfo-NHS-SS-biotin (a final concentration of 0.25 mg/mL) at 4°C for 30 min. Ultrafiltration (100-kDa cutoff) was used instead of ultracentrifugation for washing procedure. The biotin-labeled samples were washed by ice-cold PBS for 4 times (accounting for a total dilution factor of 10000) and concentrated the solution down to 50 μL. Then, 500 μL lysis buffer was added to the concentrated samples, incubated at 4°C for 30 min, and then subjected to 250 μL avidin bead slurry with gentle mixing by a horizontal rotator at 4°C for 30 min by a horizontal rotator. After 4 times washing, the EV surface proteins were eluted by 200 μL elution buffer. Samples were dried by a speedvac, resuspended in 40 μL Laemmli buffer, heated and electrophoretically run 1.5 cm into a 4-12% Invitrogen B-T gel using MOPS buffer. The gels were excised, denatured, alkalinized and digested by trypsin. The eluates were desalted by C18 solid phase extraction before nanoLC-MS/MS analysis. Three pooled (whole) EV samples were analyzed as the comparator. To mitigate technical variables, all proteins were normalized against intra-sample CD9, a surface EV marker with known equal expression between group. Surface proteins were filtered by Gene Ontology-Cellular Component of plasma membrane, and a cut-off fold-change >1 comparing between surface EV fractions and pooled (whole) EV specimens. Protein signatures based on top10% increased and decreased protein fold-changes were applied for functional prediction.

### MicroRNA-sequencing

NEBNext Small RNA Sample Library Preparation kit (NEB, Ipswich, MA) was used to construct the library with a modification for precise library size selection with high sensitivity (patent pending in the US (USSN 16/469,242) and in Europe (17881276.4)). Briefly, after 15 cycles of final PCR, the libraries with unique indices were first equal-10 μl pooled, cleaned up and mixed with custom-designed DNA ladder that contains 135 and 146 bp DNA. This size range corresponds to miRNA library with 16-27 nt insert that covers all miRNAs. After high resolution agarose gel electrophoresis, the library pool ranging from 135 to 146 bp including the DNA marker were gel purified and quantified by NEBNext Library Quant kit (NEB) using QuantStudio 5 Real-Time PCR System (Thermo Fisher, Waltham, MA). The first sequencing was performed on NextSeq 2000 sequencer (Illumina, San Diego, CA) to generate a few million reads to quantify the relative concentration of each library. The volume of each library was then adjusted to generate >3M reads per sample in the second sequencing for final data analysis. The raw fastq files were processed by Illumina Sequence Hub-miRNA analysis: trim adapter using cutadapt, map trimmed reads on miRNA precursors using SHiMPS aligner, and counts reads associated to mature miRNAs.^38^ A strict cut-off of >100 counts was applied to filter miRNAs with potential physiologic relevance.^39,40^ Log2 transform data was used for differential expression analysis and functional prediction.

### Bioinformatics and data analysis

Differential expression analysis, principal component analysis and heatmap with clustering were performed by R packages. Protein-protein interaction was analyzed by STRING (https://string-db.org). Functional/pathway enrichment analysis was performed by EnrichR (https://maayanlab.cloud/Enrichr). Functional enrichment analysis with network ensembled topology was performed by SVVATH-D (https://github.com/schuti/SVVATH-D). Prediction of miRNA gene targets was performed by miRNET (https://www.mirnet.ca). Predicted function was considered valid if adjusted p-value<0.05.

## RESULTS AND DISCUSSION

A combination of stepwise centrifugation, ultrafiltration and qEV-size exclusion chromatography (SC-UF-qEV) has been successfully employed in our previous studies to isolate EVs from the cell culture media and human plasma, resulting in an acceptable recovery yield and high specificity.^34-36^ This combined method has never been applied to human milk. Noted that human milk matrix is more complex than cell culture media and plasma, containing numerous milk fat and casein micelles which could interfere downstream EV analysis. We therefore optimized our workflow by adding low-speed centrifugation to remove milk fat, 1% w/v sodium citrate, a well-known calcium chelator,^41^ to disrupt casein micelles as the early part of SC-UF-qEV method **(Figure 3A)**. Following the isolation procedure, we validated the presence of HMEVs in the isolates following the International Society for Extracellular Vesicles (ISEV) guidelines.^42^ Nanoparticle Tracking Analysis (NTA) was performed to determine the size distribution and concentration of the isolated HMEV particles. As shown in **Figure 1B**, the majority of particles in the HMEV isolate had the diameters <200 nm, thus consistent with the small EV subpopulation as defined by the ISEV.^42^ Next, we performed transmission electron microscopy (TEM) with negative staining to confirmed the cup-shaped nano-scaled vesicle morphology **(Figure 1C)**. Finally, Western immunoblot was performed to detect three common EV markers; CD9, a surface tetraspanin, ALIX, a membrane associated protein that involved in EV biogenesis; and HSP70, a cytosolic protein commonly presented in EVs, enriched in the HMEV isolate, but not human milk (HM), defatted HM (dHM) or microvesicles (MVs) **(Figure 1D)**.

**Figure 1.**
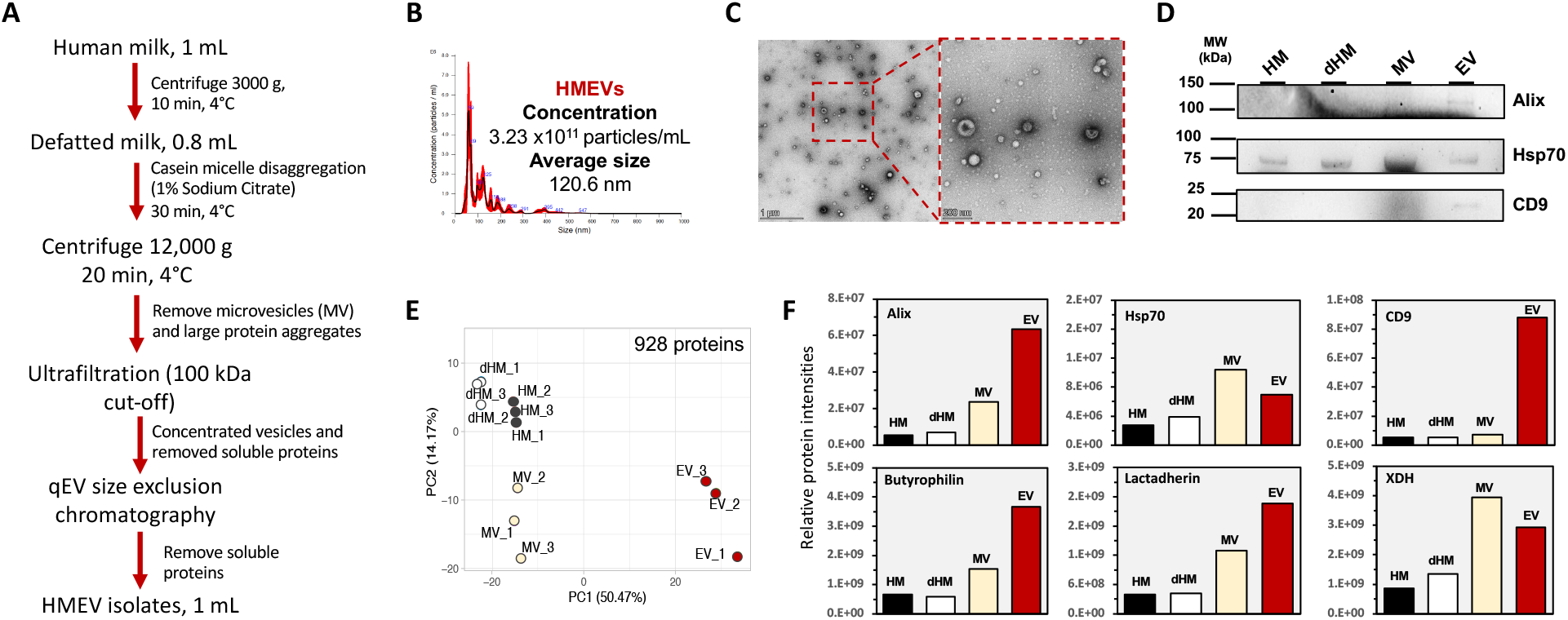
HMEV isolation and characterization. **A**. The optimized HMEV isolation workflow. B. Nanoparticle tracking analysis (NTA). **C**. Transmission electron microscope (TEM). Scale bars, 1000 nm (left panel) and 200 nm (right panel), respectively. **D**. Western immunoblotting against three common EV markers, i.e., Alix, Hsp70 and CD9. **E**. Proteomics with PCA clearly separated HMEVs from HM, dHM and MVs. **F**. Relative protein intensities of three common EV markers and HMEV-specific markers^44,45^ including butyrophilin, lactadherin and XDH. dHM, defatted HM; Hsp70, heat shock protein 70; MV, microvesicle; PCA, principal component analysis; XDH, xanthine dehydrogenase.

We previously proposed that proteomics with unsupervised machine learning, i.e., principal component analysis (PCA), could be used as part of EV quality control.^43^ To further differentiate the HMEVs from HM, defatted HM (dHM), and microvesicles (MVs), we performed a Principal Component Analysis (PCA) based on 928 proteins detected in HMEVs, HM, dHM and MVs. As seen in Figure 1D, the PCA results effectively separated the HMEVs from the other sample types. HM and dHM were more similar. MVs, or large EV subpopulation with a typical diameter of 200-1000 nm,^42^ were different from both HMEVs, HM and dHM. Targeted analysis of selected proteins demonstrated the enrichment of three EV markers, as well as lactadherin, butyrophilin and xanthine dehydrogenase (XDH), HMEV-enriched markers proposed previously,^44,45^ in HMEVs but not other milk fractions **(Figure 1F)**, thus providing an explanation to the PCA result. Taken together, our findings strongly support the use of this optimized workflow for further HMEV studies.

Through collaboration with the IMPRINT birth cohort, we sorted out 9 mothers with a history of SARS-CoV-2 infection during pregnancy (the prenatal SARS-CoV-2 group) and 9 healthy mothers (the control group) to retrieve milk samples collected at 2 weeks of lactation for HMEV isolation. Maternal demographic data and HMEV parameters, including particle numbers, protein and RNA contents, were not statistically different between groups (**Table 1**). The prenatal SARS-CoV-2 group had the COVID-19 test positive approximately 3 months (range 1.5-6 months) before delivery. Noted that both groups had not received COVID-19 vaccine before milk collection. Therefore, analyzing HMEVs collected at 2 week of lactation open up an opportunity to evidence long-term influences of post-acute SARS-CoV-2 infection upon mammary glands and lactation physiology.

**Table 1.** Maternal and human milk characteristics.

|  | **Prenatal SARS-CoV-2**<br>**(n=9)** | **Controls**<br>**(n=9)** | ***p*-value** |
| --- | --- | --- | --- |
| **Maternal variables** |  |  |  |
| **Age** (yr),<br>mean±SD | 32.8±4.4 | 30.7±4.1 | 0.29 |
| **BMI** (kg/m<sup>2</sup>),<br>mean±SD | 27.9±5.8 | 24.1±3.4 | 0.10 |
| **Parity** (n),<br>median [IQR] | 3 [2, 3] | 2 [2, 6] | 0.35 |
| **Time from COVID-19 test**<br>**positive to delivery** (week),<br>median [min-max] | 13 [6-25] | — | — |
| **Human milk, 1 mL** |  |  |  |
| **Protein concentration**<br>(after fat removal) (mg/mL) | 16.9±3.5 | 16.3±2.4 | 0.70 |
| **Total EV particle number**<br>(10<sup>11</sup> particles), median [IQR] | 6.47<br>[4.84, 8.17] | 4.75<br>[3.00, 5.71] | 0.21 |
| **Total EV protein** (µg),<br>median [IQR] | 214<br>[148, 235] | 171<br>[134, 190] | 0.30 |
| **Total EV RNA** (ng),<br>median [IQR] | 124<br>[95, 193] | 208<br>[57, 352] | 0.18 |

Abbreviations: BMI, body mass index; EV, extracellular vesicles; IQR, interquartile range.

The HMEVs isolated from 9 prenatal-SARS-CoV-2 and 9 control mothers were analyzed by proteomics, surfaceomics and miR-seq (**Figure 2**). Noted that individual samples were subjected to proteomics and miR-seq, while pooled HMEVs (3 individuals per pool) were required for surfaceomics to make 200 μg total EV proteins before surface EV protein enrichment. As a result, a total of 1,189 proteins (823 intravesicular, 366 surface), and 232 high-mid abundance miRs (count >100) were identified and quantified in HMEVs. As the quality control, targeted analysis of the common EV markers (CD9, Alix, Hsp70) and HMEV-specific markers (lactadherin, butyrophilin, XDH) as well as angiotensin converting enzyme 2 (ACE2; a known receptor of SARS-CoV2), were comparable between the prenatal-SARS-CoV-2 and control groups (**Supplementary figure 1A**).

**Figure 2.**
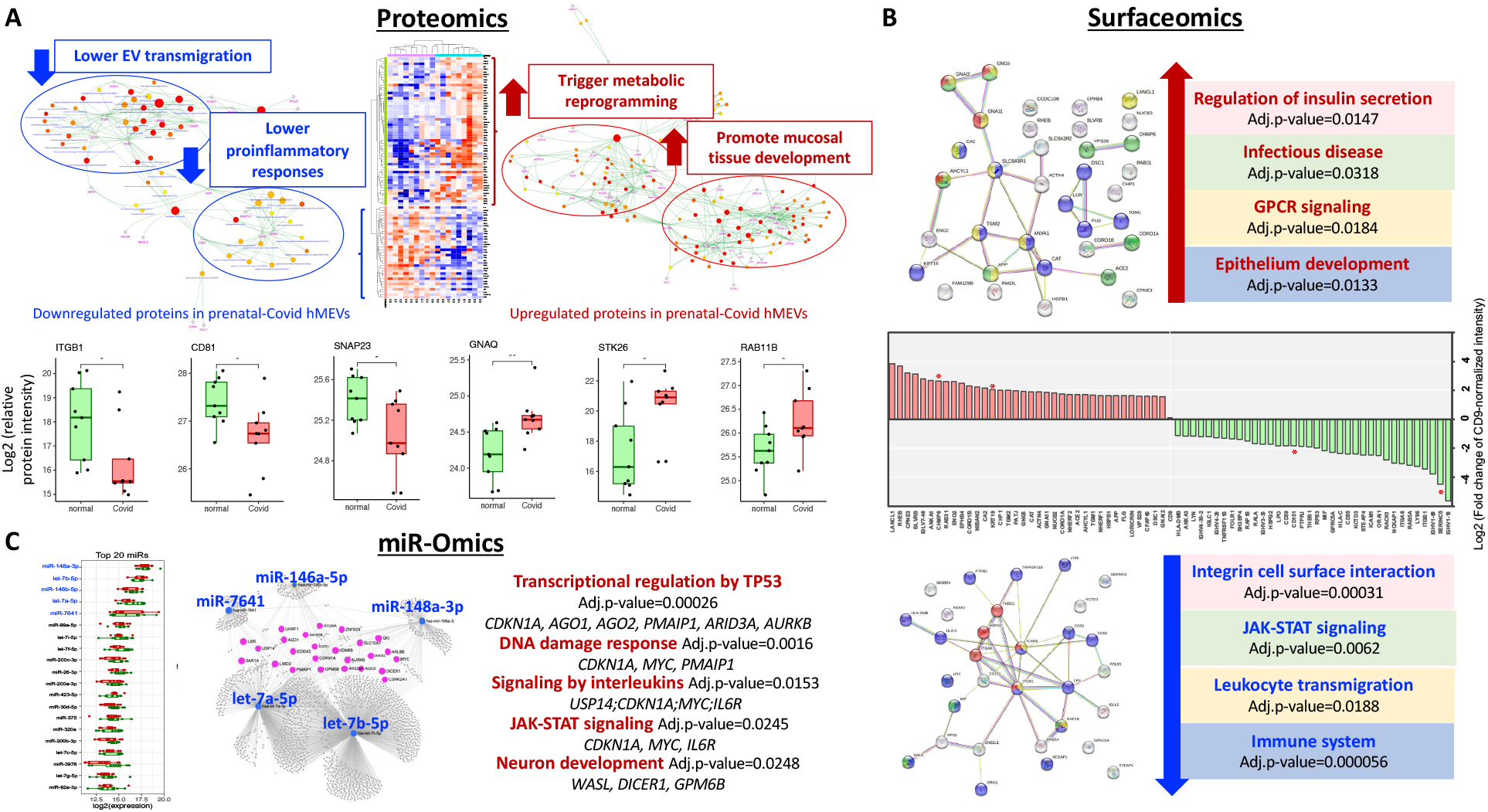
Multi-Omic analysis revealed prenatal SARS-CoV-2 infection affected HMEV molecular cargos in human milk at 2 weeks of lactation period. **A**. HMEV proteomics. Of 1143 identified proteins, 91 significantly altered proteins were analyzed by heatmap with unsupervised clustering. Functional enrichment analysis with network ensembled topology of up- and down-regulated proteins identified four functional-protein clusters. Representative six relevant EV proteins were showed as boxplots (all 30 relevant EV proteins showed in **Supplementary figure 1**). **B**. HMEV surfaceomics. Of 366 surface EV proteins, top 10% increased (36) and decreased (36) proteins were analyzed by STRING protein-protein interaction with functional enrichment analysis. Color codes represent proteins involved in predicted functions. **C**. HMEV miR-Omics. Of 2588 identified miRs, the top 20 high abundant miRs in HMEVs were illustrated. Gene target prediction with network analysis showed 24 common gene targets shared among top 5 HMEV-miRs Analyzing HMEVs at different molecular levels provided a consistent functional prediction that prenatal SARS-CoV-2 infection altered HMEV functions to promote mucosal epithelial development, mitigate inflammation, and lower systemic bioavailability, thus boosting mucosal-site specific effects. *, p<0.05; **, p<0.01.

Following differential protein expression analysis, 91 significantly altered proteins (56 upregulated and 35 downregulated; p-value <0.05) were analyzed by heatmap with unsupervised clustering (**Figure 2A**). Functional/pathway enrichment analysis with network ensemble topology demonstrated four major functional-protein domains (**Figure 2A**), corresponding to 30 HMEV proteins that may trigger functional changes in recipient cells following EV cargo transfer, specifically: (i) *Enhanced metabolic reprogramming*: upregulations of G-protein subunit alpha q (GNAQ) ^46^ and cytosolic enzymes involved in Acetyl-CoA production (ACLY, ACSS2), carbohydrate (GPI, PYGB), lipid (FASN, OLAH), purine (ADSL, PNP) and folate (MTFHD1) metabolism (**Supplementary figure 1B**); (ii) *Enhanced mucosal epithelium development*: upregulation of serine-threonine kinase STK26 mediated apical membrane remodeling,^47^, RAB11b mediated microvilli formation and intestinal cell polarity^48^ and ARHGDIB, CFL1, EPS8L2, FSCN1, PLS1, PIP, POTEJ, TWF2 that play roles in cell growth and development (**Supplementary figure 1C**); (iii) *Proinflammatory response mitigation*: upregulation of TPT1 mediated T and NK cell suppression^49^ and USP14 stabilizing IDO1 immunosuppressive molecule,^50^ and downregulation of SNAP23 regulated cytokine release,^51^ CD81 mediated T_h_2 polarization^52^ and MATN2/3 extracellular matrix proteins involved in proinflammatory responses^53,54^ (**Supplementary figure 1D**); (iv) *Lowered systemic bioavailability*: downregulation of surface adhesion molecules ITGB1/4 and LAMB1 may reduce HMEV migration and translocation through integrin-laminin interaction^55-57^ (**Supplementary figure 1E**).

Surfaceomics revealed 366 EV surface proteins successfully enriched by the biotin-avidin bead capture. Since surfaceomics was performed using the pooled samples (3 samples generated one pooled EVs; 3 pooled samples/group) which lacked of statistical power, functional interpretation was then performed using protein fold-change signatures (Top10% increased and decreased proteins; n=36 each) with STRING protein-protein interaction with functional enrichment analysis (**Figure 2B**). Accordingly, functional prediction revealed HMEVs of the prenatal SARS-CoV-2 group may promote metabolic regulation through insulin secretion, G protein-couple receptor (GPCR) signaling, and epithelial development, and decrease functions in the domains of integrin cell surface interaction, JAK-STAT signaling, leukocyte transmigration and immune response (**Figure 2B**).

A total of 2588 miRs with at least one count were identified across 18 individual HMEV samples. Nonetheless, only 232 miRs passed a strict cut-off of >100 counts, representing high-mid abundance HMEV-miRs with potential physiologic relevance.^39,40^ Interestingly, differential expression analysis showed no significantly altered miRs between the prenatal SARS-CoV-2 and control groups (**Supplementary figure 2**), suggesting a tightly regulated mechanism of miR packaging of mammary epithelial cells to preserve HMEV essential functions. In this direction, the top 5 highest abundance HMEV-miRs, i.e., miR-148a-3p, let7b-5p, let7a-5p, miR146a-5p and miR-7641 were further annotated for their gene targets using miRNET. Functional enrichment analysis was performed on the shared targeted genes among top miRs. As a result, **Figure 2C** showed that top 5 highest abundance HMEV-miRs may exhibit physiologic functions involving transcriptional regulation, cell growth and development, and immunomodulation. Noted that miR-148a is the most abundance HMEV-derived microRNA,^11,58^ influencing various functions, including DNA methylation through DNA methyltransferase 1,^59^ cell and tissue development,^60^ inflammation,^61,62^ and neuroprotection.^63,64^ Also, miR-146a-5p is known to regulate type I interferon signaling^65,66^ and involved in cellular responses upon viral infection.^67,68^

Taken together, our findings demonstrate that prenatal SARS-CoV-2 infection (1-6 months before delivery) does have post-acute infection effects on HMEV-derived proteins, but not miRNAs, at 2-week of lactation. While physiologic relevance of HMEV-miRs are preserved, downregulation of surface adhesion molecules (**Figure 2A**,**B** and **Supplementary figure 1E**) have potential to change tissue distribution and systemic bioavailability of ingested HMEVs.^14-17^ In addition, upregulation of proteins involving metabolic regulation, energy production and mucosal tissue development (**Figure 2A**,**B** and **Supplementary figure 1B-D**) can be seen as the enhanced mucosal-site specific functionalities of HMEVs following prenatal SARS-CoV-2 infection. thereby offering protection of breastfed infants against intestinal and respiratory diseases. HMEVs had been shown to exhibit broad antiviral effects in vitro (i.e., HIV, rotavirus, RSV, CMV, Zika).^69-72^ A possible hypothesis is that HMEVs may activate the innate defense within recipient cells particularly type I interferon, through miR-146a-5p^65-68^ (**Figure 2C**). In this direction, HMEVs from prenatal SARS-CoV-2 mothers may boost mucosal-site specific functions to prevent respiratory viral infections, including SARS-CoV-2, in breastfed infants.

In conclusion, this study communicated, for the first time, that prenatal SARS-CoV-2 infection affected HMEV molecular cargos in a way that promote mucosal-site specific functions of HMEVs. Nonetheless, these preliminary findings require further validation in a larger sample size and follow-up across multiple stages of lactation to determine whether HMEV alterations after prenatal SARS-CoV-2 infection are temporary or persistent. The effect of maternal COVID-19 vaccination and of exposure to both SARS-CoV-2 infection and vaccination on HMEV molecular cargo also remains to be determined. Direct evidence of HMEV functions at cellular levels is lacking, including the HMEV molecular-function relationship on tissue development, immunomodulation and viral inhibition. Addressing these issues have potentials to deliver scientific evidence that inform public health strategies and breastfeeding practice to improve child health outcomes in the post-COVID era.

## ACKNOWLEDGMENTS

We thank all mothers who participated in the project.

## AUTHOR CONTRIBUTIONS

Conceptualization: S.C. and A.L.M.; Methodology, S.C., X.Z., D.B., S.L., D.S.N., K.D.G., M.A.S. and A.L.M.; Software, S.C., M.W.; Validation, S.C., X.Z., D.B., S.L., L.N.R., D.S.N., K.D.G., M.A.S. and A.L.M.; Formal Analysis, S.C., H.C., X.Z., D.K., D.B., W.H., M.W.; Investigation, S.C., H.C., X.Z., D.K., D.B., W.H., M.W.; Resources, X.Z., S.L., L.N.R., D.S.N., K.D.G., M.A.S. and A.L.M.; Data Curation, R.P., S.C.C., and A.R.B.; Writing – Original Draft Preparation, S.C.; Writing – Review & Editing, J H.C., X.Z., D.K., D.B., W.H., M.W., R.P., S.C.C., A.R.B, S.L., L.N.R., D.S.N., K.D.G., M.A.S. and A.L.M.; Visualization, S.C., H.C., D.K., D.B.; Supervision, D.S.N., K.D.G., M.A.S. and A.L.M.; Project Administration, R.P., S.C.C., A.R.B.; Funding Acquisition, X.Z., K.D.G., M.A.S. and A.L.M. All authors have read and agreed to the published version of the manuscript.

## SUPPLEMENTARY MATERIALS

**Supplementary figure 1**. Targeted protein analysis of HMEV proteome acquired from 9 prenatal SARS-CoV-2 vs. 9 control mothers. **A**. EV markers and ACE2. **B**. Protein involved in metabolic reprogramming. **C**. Proteins involved in mucosal tissue development. **D**. Proteins involved in pro-inflammatory responses. **E**. Surface adhesion molecules. ns, not significant. *, p<0.05. **, p<0.01.

**Supplementary figure 2**. HMEV miR analysis. **A**. PCA based on all 2,588 identified miRs. Total 232 miRs passed a cutoff of 100 counts. **B**. Mean-variance trend plot for quality control. **C**. Volcano plot show no differentially expressed miRs between groups.

