## Supplementary figures and images for "Prenatal SARS-CoV-2 infection alters postpartum human milk-derived extracellular vesicles"

### Supplementary figures

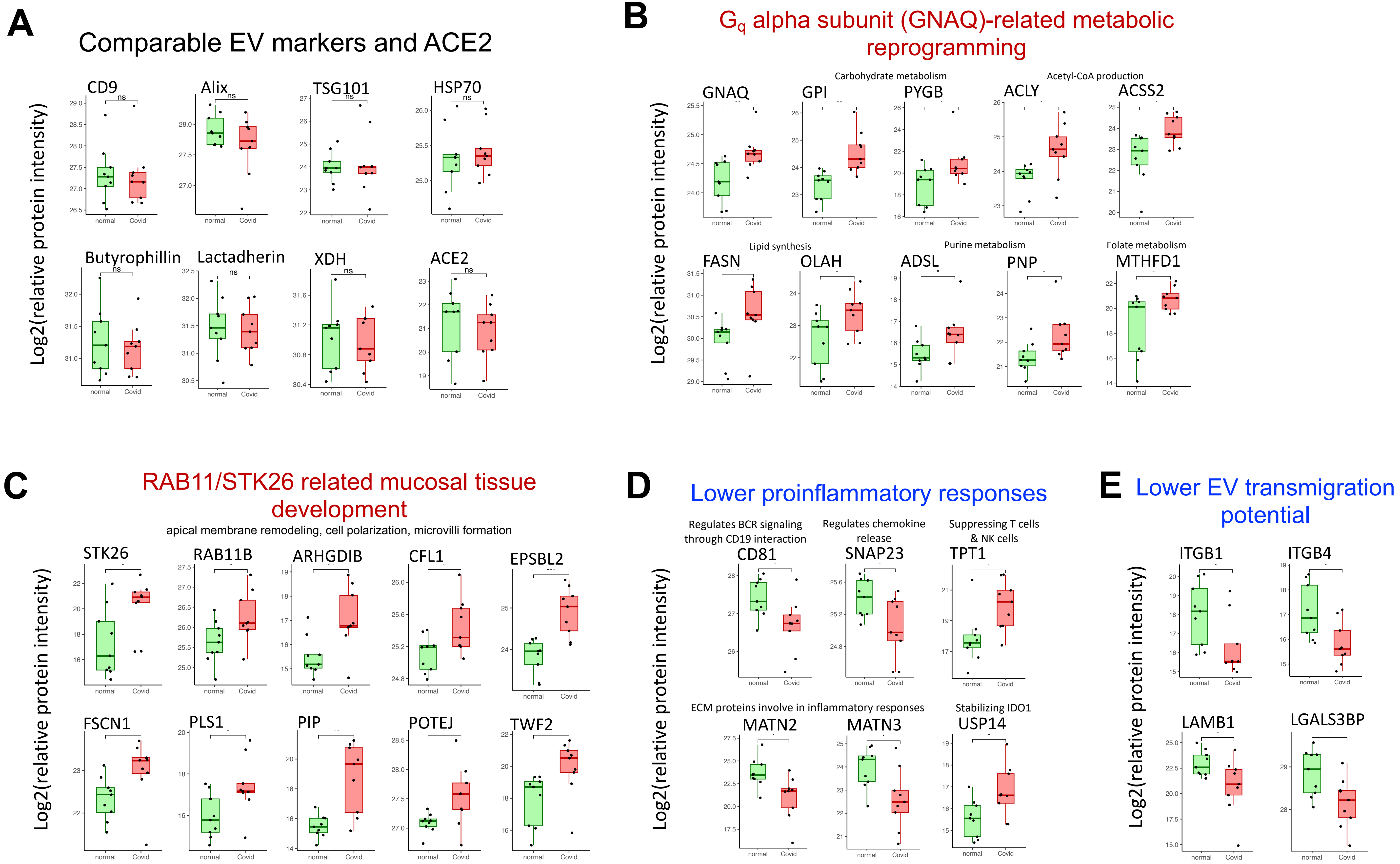

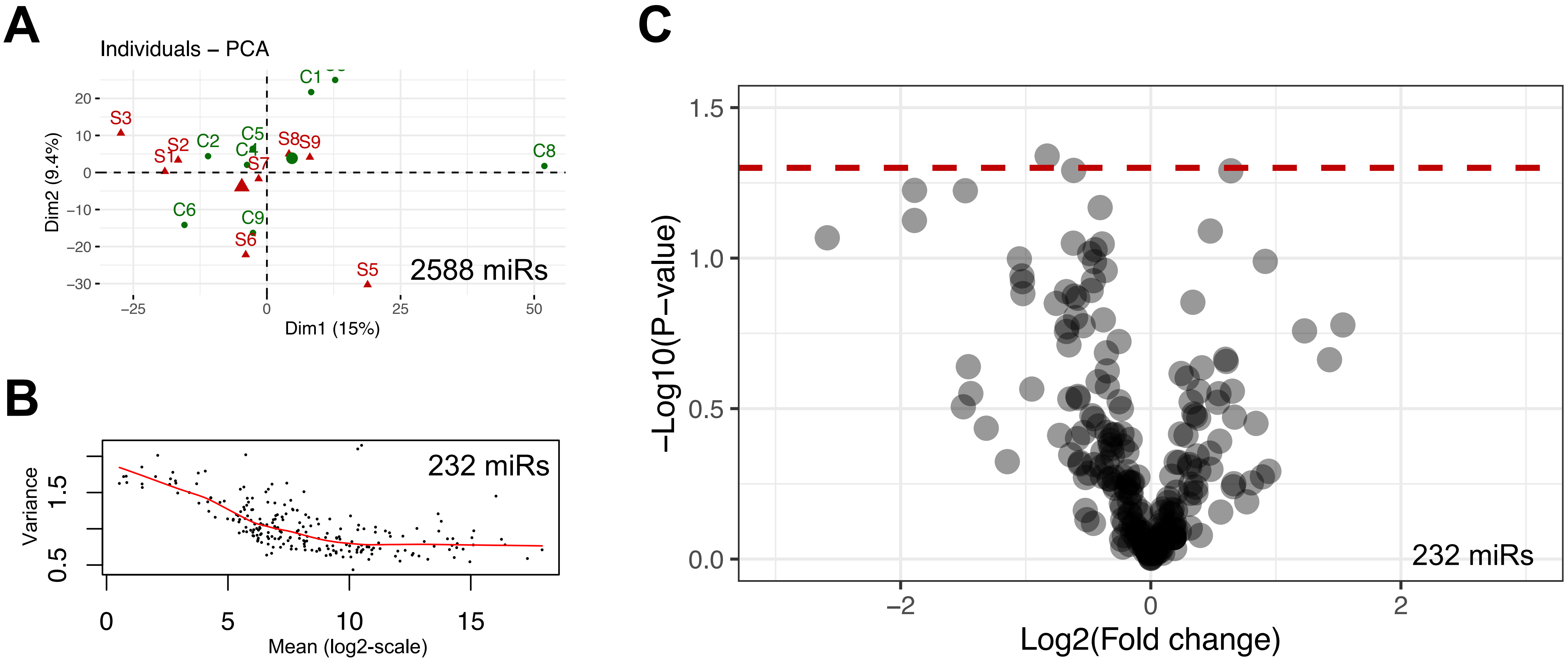
